# The Endoplasmic Reticulum Proteostasis Regulator ATF6 is Essential for Human Cone Photoreceptor Development

**DOI:** 10.1101/2020.10.04.325019

**Authors:** Heike Kroeger, Julia M. D. Grandjean, Wei-Chieh Jerry Chiang, Daphne Bindels, Rebecca Mastey, Jennifer Okalova, Amanda Nguyen, Evan T. Powers, Jeffery W. Kelly, Neil J. Grimsey, Michel Michaelides, Joseph Carroll, R. Luke Wiseman, Jonathan H. Lin

## Abstract

Dysregulation of the endoplasmic reticulum (ER) Unfolded Protein Response (UPR) is implicated in the pathology of many human diseases associated with ER stress. Inactivating genetic variants in the UPR regulator *Activating Transcription Factor 6 (ATF6)* cause severe congenital heritable vision loss in patients by an unknown pathomechanism. To investigate this, we generated retinal organoids from patient iPSCs carrying *ATF6* disease-causing variants and *ATF6* null hESCs generated by CRISPR. Interestingly, we found that cone photoreceptor cells in *ATF6* mutant retinal organoids lacked inner and outer segments concomitant with absence of cone phototransduction gene expression; while rod photoreceptors developed normally. Adaptive optics retinal imaging of patients with disease-causing variants in *ATF6* also showed absence of cone inner/outer segment structures but preserved rod structures, mirroring the phenotypes observed in our retinal organoids. These results reveal that *ATF6* is essential for the formation of human cone photoreceptors, and associated absence of cone phototransduction components explains the severe visual impairment in patients with *ATF6* -associated retinopathy. Moreover, we show that a selective small molecule ATF6 activator compound restores the transcriptional activity of *ATF6* disease-causing variants and stimulates the growth of cone photoreceptors in patient retinal organoids, demonstrating that pharmacologic targeting of ATF6 signaling is a therapeutic strategy that needs to be further explored for blinding retinal diseases.

## Results and Discussion

*Activating Transcription Factor 6 (ATF6)* encodes an ER resident type 2 transmembrane protein that controls a key signal transduction pathway of the mammalian Unfolded Protein Response (UPR) ^1,2^. In response to pathologic or physiologic events that disrupt ER homeostasis, ATF6 migrates from the ER to the Golgi, where proteases cleave the protein to release the cytosolic, N-terminal ATF6 transcriptional activator fragment ^3^. The liberated ATF6 transcription factor enters the nucleus to induce ER chaperones and protein folding enzymes that increase the biosynthetic capacity of the ER, allowing the cell to adapt to and survive episodes of ER stress ^4,5^.

In people, disease-causing variants in *ATF6* are a cause of heritable vision loss retinal diseases, primarily achromatopsia, and to a lesser extent cone-rod dystrophy^6,7^. Puzzlingly, ATF6 shares no biologic similarities with any other genes that cause achromatopsia - *CNGB3, CNGA3, PDE6C, PDE6H*, and *GNAT2* – all of which mediate cone phototransduction^6^. We previously demonstrated that *ATF6* variants impair its transcriptional activity by interrupting essential steps in the ATF6 signal transduction pathway ^8^. For example, a tyrosine to asparagine conversion at residue 567 (Y567N) in the ATF6 luminal domain impeded ER-to-Golgi trafficking of the full-length ATF6; while an arginine to cysteine conversion at residue 324 (R324C) in the ATF6 bZIP domain prevented DNA binding by the ATF6 transcriptional activator fragment ^8^. Thus, patients homozygous for *ATF6* disease alleles (*ATF6*^*hom*^) generate transcriptionally incompetent ATF6 proteins and have severely impaired vision from birth; while heterozygous (*ATF6*^*het*^) carriers express a normal copy of *ATF6* and have normal vision ^6,8^. To determine why *ATF6* variants cause vision loss, we generated induced pluripotent stem cells (iPSCs) from fibroblasts of patients with achromatopsia who were homozygous carriers of *ATF6* disease-causing variants ^6,8^. For comparison, we also generated iPSCs from fibroblasts of unaffected family members who were heterozygous carriers of these variants. We then differentiated these *ATF6*^*hom*^ and *ATF6*^*het*^ iPSCs into retinal organoids following established protocols and examined their photoreceptors ^9^.

### *ATF6* Mutant Retinal Organoids Have No Cone Structures

In the retina, cone and rod photoreceptors are morphologically distinguishable by the eponymous shapes of their polarized inner/outer segments ^10-12^. The photoreceptor inner segment begins where the photoreceptor cell soma protrudes beyond the external limiting membrane of the retina and contains abundant biosynthetic (ER/golgi) and metabolic (mitochondria) organelles; more distally, the inner segment connects with the outer segment, a specialized sensory cilia devoted to light detection, that houses hundreds of membranous discs containing visual pigments and phototransduction proteins. In mature human cones, the entire inner/outer segment adopts a rotund ovoid morphology beginning at the external limiting membrane that rapidly tapers off distally (Fig. 1d, schematic illustration). By contrast, in rods, the inner/outer segment adopts a slender cylindrical morphology throughout its length (Fig. 1d, schematic illustration).

**Fig. 1.**
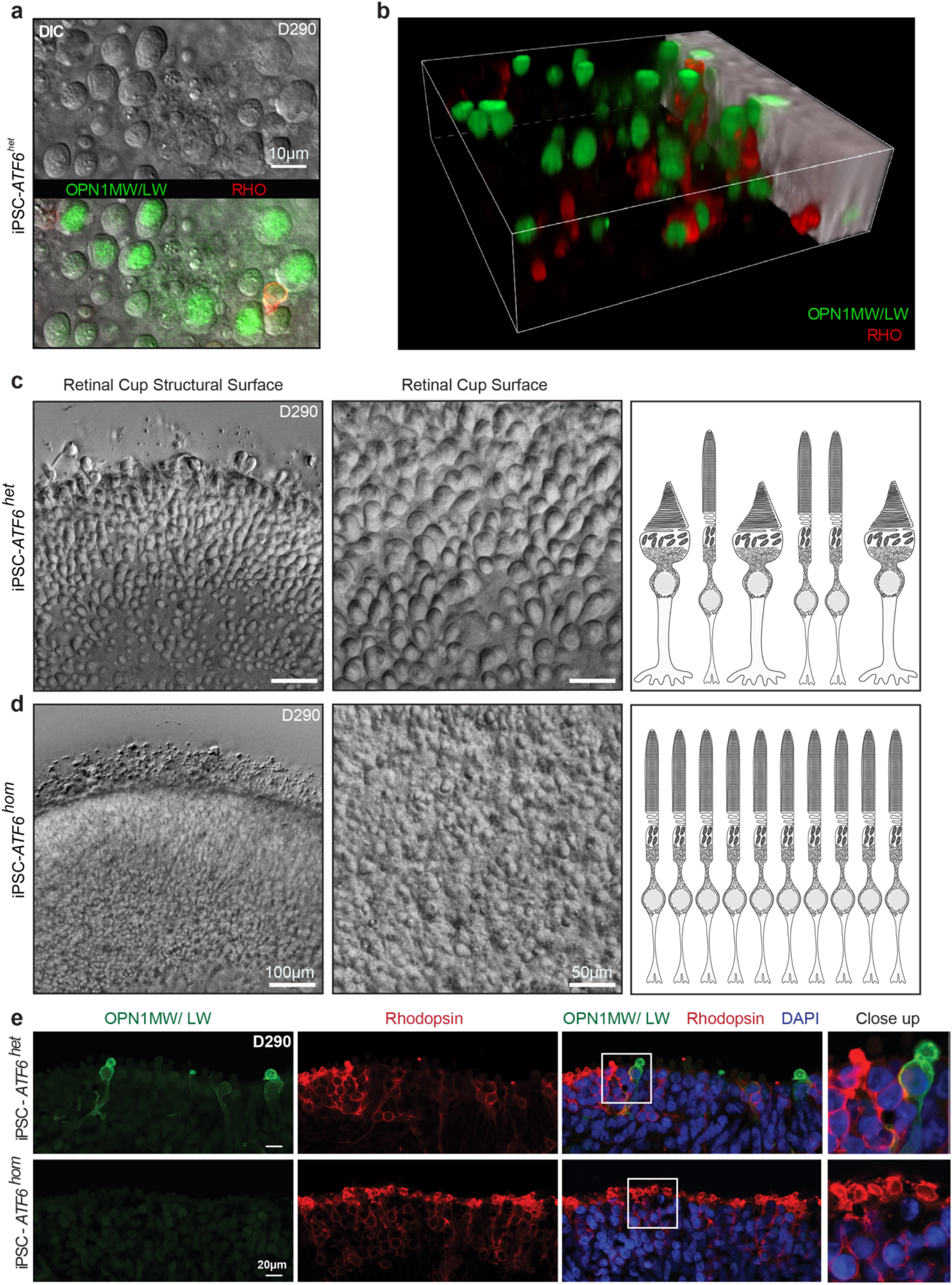
*ATF6* mutant retinal organoids lack cone inner/outer segment structures. **(a)** Top-down view of DIC microscopy Z-stack series of rhodopsin (RHO) and red/green cone opsin (OPN1MW/LW) labeled retinal organoids shows large round structures on the surface of retinal organoids. Superimposed immunofluorescence microscopy shows cone opsin OPN1MW/LW labeling (green) colocalizing with round structures and exclusion of rhodopsin labeling (red). **(b)** Snapshot from 3-D reconstruction video (Supplementary Video 1) of confocal fluorescent microscopy images show that round structures on retinal organoid surfaces adopt ovoid morphologies in 3-D encapsulating OPN1MW/LW (green) and excluding rhodopsin (red) consistent with rudimentary cone inner/outer segments. **(c)** Images were taken at the same optical magnification, digital zoom was used to demonstrate cellular details of the retinal organoid surface. Overview (left panel) and enlargement for detailed view (middle panel) DIC microscopy images show abundant cone structures on surface of *ATF6*^*het*^ retinal organoids. **(d)** *ATF6*^*hom*^ retinal organoids show absence of cone structures with retention of rod structures. Right panel cartoons in (c) and (d) summarize absence of cone structures in *ATF6*^*hom*^ retinal organoids. **(e)** Confocal fluorescence microscopy images show absence of OPN1MW/LW (green) and preservation of rhodopsin (red) labeling in 290-days (D290) old retinal organoids differentiated from homozygous *ATF6* disease mutation iPSCs (iPSC-*ATF6*^*hom*^) (bottom rows) compared to heterozygous *ATF6* iPSCs (iPSC-*ATF6*^*het*^) (top rows). DAPI (blue) identifies nuclei. Three independent experimental repeats were performed (n=3).

Maturing photoreceptors in retinal organoids recapitulate morphologic and molecular features of cone and rod photoreceptors ^13-15^. With prolonged *in vitro* culturing, round ovoid protrusions emerged from the surfaces of *ATF6*^*het*^ retinal organoids that were morphologically consistent with nascent cone inner/outer segments and contained cone opsin proteins (**Fig. 1a, 1b, Supplementary Video 1**). By contrast, rhodopsin was excluded from these ovoid protrusions and confined to slimmer, cylindrical structures consistent with rod inner/outer segments. The presence of ovoid or cylindrical protrusions on the retinal organoid surface enabled rapid visual identification and longitudinal tracking of developing cones and rods by live organoid imaging. In doing so, we discovered a striking and unexpected malformation of cones in retinal organoids lacking functional ATF6.

By 200 to ∼300-days of differentiation, we saw abundant cone structures on retinal organoid surfaces of *ATF6*^*het*^ iPSCs generated from family members with normal vision (**Fig. 1c, Supplementary Video 2**). *ATF6*^*hom*^ iPSCs also differentiated and formed retinal organoids that appeared grossly identical to *ATF6*^*het*^ retinal organoids (**Fig. S1**). However, microscopic examination of the surfaces of *ATF6*^*hom*^ patient retinal organoids showed no ovoid cone structures and instead revealed smoother contours consistent with fine packing of slender rods (**Fig. 1d, Supplementary Video 3**). In keeping with these live retinal organoid surface imaging findings, rod and cone inner/outer segments could be detected and distinguished by non-overlapping expression of rhodopsin and red/green cone opsin proteins (OPN1MW/ LW) on immunofluorescent confocal microscopy of fixed cross-sections of 290-day old *ATF6*^*het*^ retinal organoids, (**Fig. 1e, top**). However, red/green cone opsin expression was completely abolished in mutant retinal organoids, though rhodopsin protein expression remained abundantly detected (**Fig 1e, bottom**). Absence of cone structures was observed in all retinal organoids homozygous for *ATF6* disease-causing variants and at all timepoints during retinal organoid differentiation and culturing *in vitro* (up to ∼300 days).

To test if the malformation phenotype observed in iPSC lines carrying human *ATF6* disease alleles arose directly via loss of *ATF6*, we next created isogenic *ATF6* null hESCs by CRISPR-mediated indel introduction into exon 1 of the human *ATF6* gene (*ATF6*^*ex1Δ/ex1Δ*^) in wild-type hESCs (**Fig. S2a**). We confirmed complete loss of ATF6 protein and downregulation of canonical downstream ATF6 transcriptional target genes in *ATF6*^*ex1Δ/ex1Δ*^ hESCs such as *GRP78/BiP* (**Fig. S2b,c**)^4,5^. When we differentiated isogenic *ATF6*^*+/+*^ and *ATF6*^*ex1Δ/ex1Δ*^ hESCs into retinal organoids, we saw abundant ovoid cone structures on wild-type organoid surfaces but no ovoid protrusions on retinal organoids lacking ATF6, similar to what we observed in patient iPSC-derived retinal organoids (**Fig. S2d,e, Supplementary Videos 4, 5**). These iPSC and hESC findings revealed an unexpected and fully penetrant cone malformation phenotype – absence of inner/outer segment structures – arising in retinal organoids with inactivation of ATF6.

### Patients with disease-causing variants in *ATF6* Do Not Have Cone Inner/Outer Segment Structures in their Retinas

To directly determine if cones in patients carrying *ATF6* disease-causing variants were malformed as those observed in retinal organoids, we performed adaptive optics scanning laser ophthalmoscopy (AOSLO) to examine the cellular shape of photoreceptors in patients carrying *ATF6* disease alleles. These included the same individuals who contributed fibroblasts for our retinal organoid studies. AOSLO enables non-invasive, imaging of the human retina at single cell resolution ^16^, and cone and rod morphologies are readily distinguished using this scanning modality ^17^. In patients with wild-type *ATF6* and normal vision, AOSLO showed numerous cone inner and outer segments in the parafovea, a specialized region of the human retina highly enriched in cones (**Fig. 2a, b**). By contrast, AOSLO imaging of *ATF6*^*hom*^ patients showed complete absence of cone inner and outer segments, but retained rod structures (**Fig. 2d, e)**. All homozygous carriers of *ATF6* disease alleles lacked cone structures with AOSLO imaging ^18^. Furthermore, longitudinal AOSLO imaging of individual *ATF6*^*hom*^ patients at follow-up evaluations (up to 3 years duration) showed that the absence of cone structures was a stationery phenotype (no episodes of growth/decay). These imaging findings in *ATF6*^*hom*^ patients revealed that the cone malformation phenotype found in retinal organoids accurately reflected the photoreceptor pathology in affected patients. Taken together, our patient imaging and retinal organoid findings identify a critical role for ATF6 in the generation of cone structures in human photoreceptors.

**Fig. 2.**
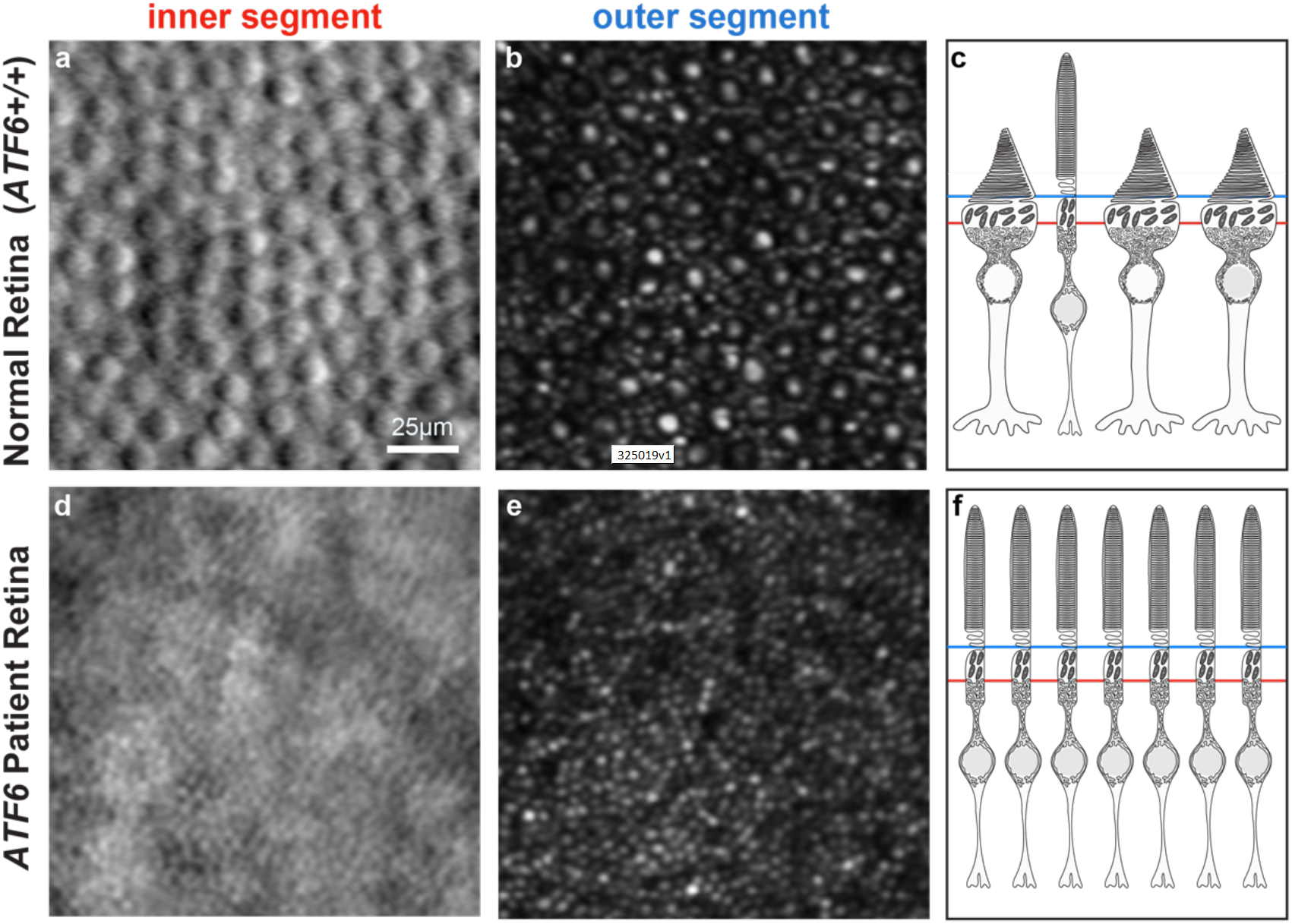
Adaptive optics retinal imaging of patients with *ATF6*^*hom*^ variants reveal absence of cone inner and outer segments. **(a, b, c)** Abundant cone inner segments (a) and outer segments (b) are identified in the retina of a normally-sighted patient (*ATF6*^*+/+*^) by split detector (a) and confocal (b) adaptive optics scanning laser ophthalmoscopy (AOSLO) (large round structures). **(d, e, f)** AOSLO of a patient with achromatopsia caused by *ATF6*^*hom*^ variants shows absence of cone inner (d) and outer (e) segments, but preservation of rod inner and outer segments. Cartoons on the right depict AOSLO planes of scanning at the levels of the inner segment (red line) and outer segment (blue line) and summarize absence of cone structures in the patient with *ATF6*^*hom*^.

### Disruption of Cone Opsin and Phototransduction Gene Expression in *ATF6* Mutant Retinal Organoids

To further investigate the impact of *ATF6* variants on cone development, we next performed RNA-seq on retinal organoids and examined transcript levels of cone-specific and rod-specific genes previously defined by human retina transcriptome profiling ^19^. The median expression of the cone gene panel in *ATF6*^*hom*^ retinal organoids was significantly reduced relative to *ATF6*^*het*^ organoids, affecting ∼75% of genes in the cone gene panel (**Fig. 3a, Supplemental Table S1**). Interestingly, within the cone gene panel, the most severely reduced genes included all cone phototransduction genes - *CNGB3, CNGA3, PDE6C, PDE6H*, and *GNAT2* (**Fig. 3b, Supplemental Table S2**). Cone visual pigment genes including red (*OPN1LW*) and green (*OPN1MW*) cone opsin also showed reduced expression in *ATF6* mutant retinal organoids (**Fig. 3c, Supplemental Table S2**), corroborating absence of the red/green cone opsins observed by fluorescent microscopy (**Fig. 1e**). By contrast, the median expression of the rod gene panel, including *rhodopsin* (*RHO*) and most rod phototransduction genes, was not significantly altered in *ATF6* mutant organoids (**Fig. 3a, 3b, Supplemental Table S2**). These findings identified a striking defect in cone genes required for light detection and phototransduction in retinal organoids lacking ATF6.

**Fig. 3.**
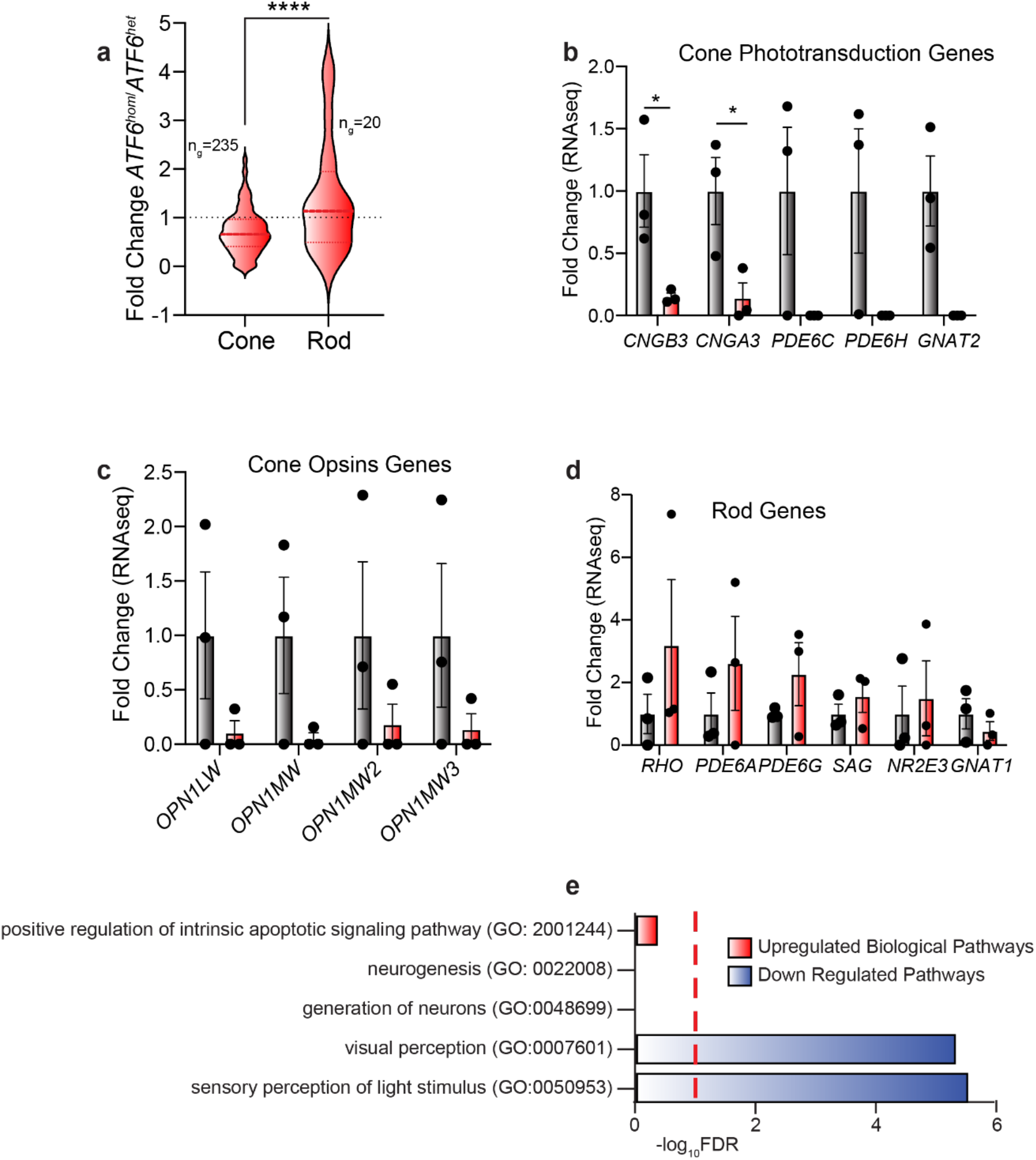
Cone visual pigments and cone phototransduction gene expression is severely reduced in ATF6 mutant retinal organoids. **(a)** RNA-seq profiling reveals significant loss of cone photoreceptor transcripts in *ATF6*^*hom*^ retinal organoids compared to *ATF6*^*het*^ retinal organoids. Violin plots show expression levels of 235 cone photoreceptor genes and 20 rod photoreceptor genes identified by RNAseq in *ATF6*^*hom*^ retinal organoids normalized to *ATF6*^*het*^ retinal organoids. The thick dashed horizontal line marks the median level of gene expression, and the thin horizontal lines delimit the upper and lower quartiles of genes in each violin plot. The complete gene sets are shown in Supplemental Table S2. ****P<0.0001, t=5.234. (**b**) Cone phototransduction genes transcripts measured by RNA-seq from *ATF6*^*hom*^ retinal organoids (red columns) are shown relative to transcript levels in *ATF6*^*het*^ retinal organoids (grey columns). Error bars represent mean +/- standard deviation, and points represent individual retinal organoids. * p<0.05. (**c**) Cone opsins (visual pigments) genes transcripts measured by RNA-seq from *ATF6*^*hom*^ retinal organoids are shown relative to transcript levels in *ATF6*^*het*^ retinal organoids. (**d**) Rod phototransduction and rhodopsin genes transcripts measured by RNAseq from *ATF6*^*hom*^ retinal organoids are shown relative to transcript levels in *ATF6*^*het*^ retinal organoids. **(e)** Gene ontology analysis identifies up-regulated (red) and down-regulated (blue) biological processes in transcriptional datasets from ATF6 mutant organoids compared to heterozygous organoids. Red-dashed line indicates false discovery rate (FDR) = 0.1. Three independent experimental repeats were performed (n=3).

Gene ontology (GO) analysis of the RNA-seq datasets independently identified that the most significantly down-regulated GO biologic processes in *ATF6* mutant retinal organoids were visual perception and light detection (**Fig. 3e, Supplemental Table 3**). By contrast, genes involved in neurogenesis and neuronal development GO biological processes were not significantly changed in *ATF6* mutant retinal organoids (**Fig. 3e, Supplemental Table 3**); but showed some reduction by targeted examination using Gene Set Enrichment Analysis (GSEA) (**Fig. S3a, b, FDR=0**.**1**). Expression of *CRX* also showed significant reduction in *ATF6* mutant retinal organoids, but other cone/rod photoreceptor cell fate genes were not significantly altered (**Fig. S3c**) ^20^. These findings suggested that, in addition to the severely negative effect on cone photodetection/transduction pathways, some early aspects of photoreceptor cell fate commitment and identity specification were also negatively impacted in *ATF6* mutant retinal organoids.

Interestingly, GO and GSEA revealed no significant change in apoptosis pathway gene expression (**Fig**.**3e, S3d, Supplemental Table 3**); furthermore, no increased expression of UPR genes implicated in ER stress-induced cell death including *CHOP/DDIT3, DR5, CKB*, and *ATF4* was found in *ATF6* mutant retinal organoids (**Fig. S3e**) ^21-25^. These results indicate that loss of *ATF6* did not trigger profound cell death in retinal organoids, consistent with no gross differences in organoid differentiation, growth, and appearance by visual inspection (**Fig. S1**).

### A Small Molecule ATF6 Agonist Rejuvenates Diseased Cone Photoreceptors

All *ATF6* disease alleles identified to date impair ATF6 signaling. Therefore, augmentation of ATF6 signaling could help patients with vision loss diseases arising from these alleles by restoring cone development. However, this strategy needs to be precisely tailored to the distinct pathomechanisms caused by different *ATF6* disease alleles ^8^. We previously identified soluble, non-toxic small molecules that activate ATF6 signaling in cell culture and mice by promoting *ATF6* reduction and monomerization (**Fig. 4a**)^26-28^. By increasing this pool of reduced *ATF6* competent to exit the ER, these compounds increase the levels of the active *ATF6* transcriptional activator fragment available for downstream signaling ^29^. This class of small molecule proteostasis regulators offers a potential chemical strategy to counter the molecular pathomechanism of Class 1 *ATF6* disease alleles arising from decreased levels of ATF6 exiting the ER. To test this strategy, we evaluated the efficacy of a lead small molecule ATF6 agonist, AA147, on a Class 1 *ATF6* variant, *Y567N*, found in patients with achromatopsia 6. Consistent with prior studies, treatment of HEK293 cells expressing the Y567N ATF6 disease-associated protein with the chemical ER stressor, DTT, failed to increase proteolytic release of the active ATF6 transcription factor (ATF6^NT^) or increase levels of the ATF6 target protein GRP78/BiP (**Fig. 4b**)^8^. By contrast, media supplementation with AA147, but not the inactive AA147 analog RP22, robustly restored production of the ATF6 transcriptional activator fragment and increased levels of the ATF6 downstream target gene, GRP78/BiP (**Fig. 4b**). These findings demonstrate that AA147 potently rescues the transcriptional function of the Y567N human ATF6 achromatopsia-associated disease mutant.

**Fig. 4.**
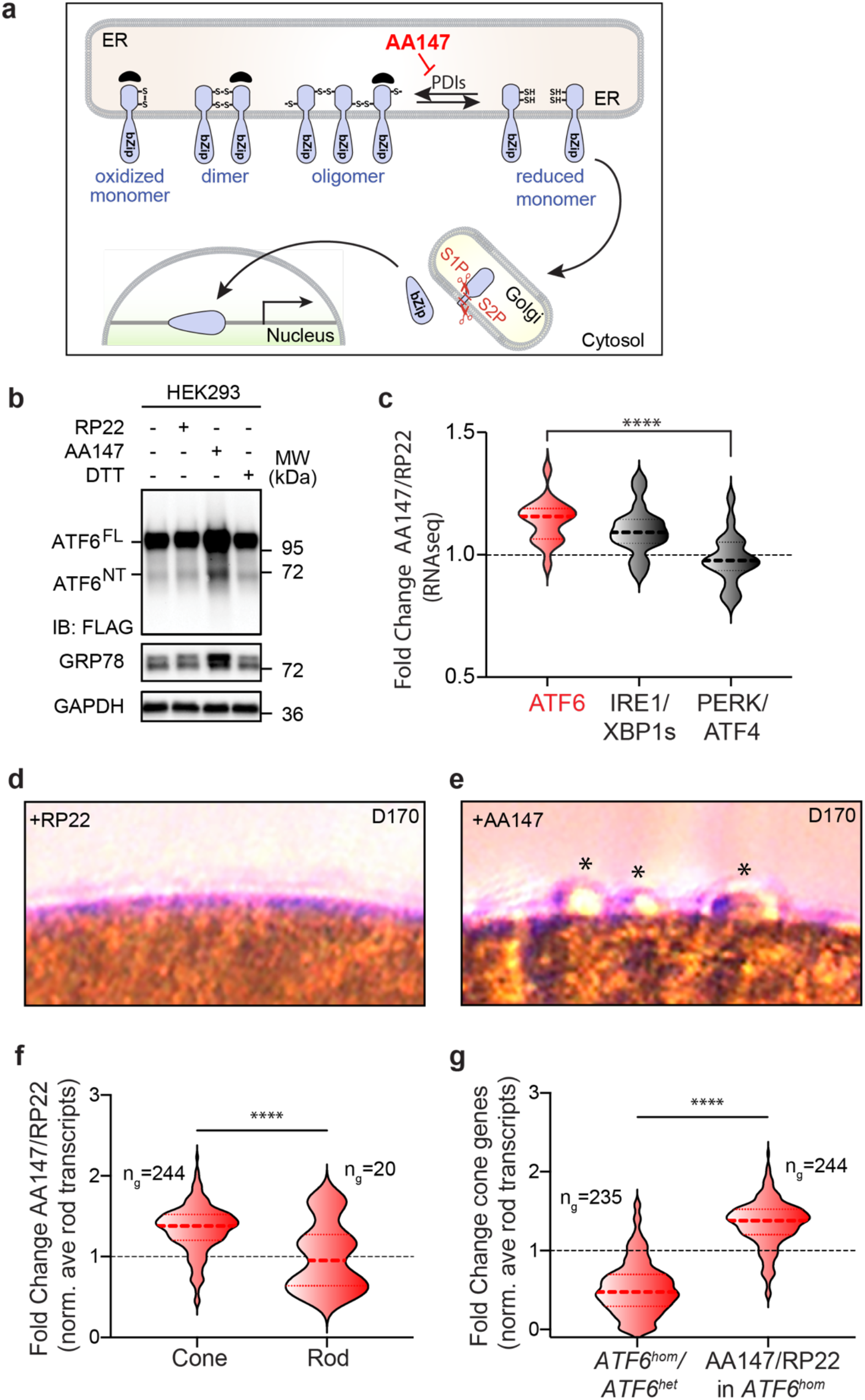
Small molecule proteostasis agonist rescues ATF6 disease mutant transcriptional activity and promotes cone development in mutant retinal organoids. **(a)** Schematic cartoon shows mechanism of ATF6 signaling pathway activation by small molecule proteostasis regulator, AA147, through increased generation of reduced ATF6 monomer in the ER by inhibiting protein disulfide isomerase (PDI). (**b)** HEK293 cells expressing FLAG-tagged ATF6 bearing the Y567N disease mutation were cultured with AA147 (10μM), RP22 (10μM), DTT (2mM), or DMSO solvent for 24h, and protein lysates were immunoblotted for FLAG, GRP78, or GAPDH (loading control). Positions of full-length ATF6 (ATF^FL^) and the ATF6 transcriptional activator amino-terminal fragment (ATF6^NT^) proteins are shown. (**c)** RNA-seq profiling of 170-day old (D170) *ATF6*^*hom*^ retinal organoids treated with AA147 or RP22 (10μM) for 50 days. Violin plots show expression levels of ATF6-, IRE1/XBP1-, and PERK/ATF4-target genes of AA147-treated relative to RP22-treated retinal organoids. The thick dashed horizontal line marks the mean; and the thin horizontal lines delimit the upper and lower quartiles of gene expression in each violin plot. The complete gene sets are shown in Supplemental Table S5. ****p<0.0001, F=9.436, RNA was generated as pooled replicates from two organoids per treatment from two independent experimental repeats n=2; data were collected using three technical repeats n=3. (**d, e)** DIC imaging shows presence of nascent cone outer segments (*) on *ATF6*^*hom*^ retinal organoid surfaces after AA147 (10μM) (e) but not RP22 (10μM) treatment (d) for 50 days. (**f)** RNA-seq profiling reveals significant gain of cone photoreceptor transcripts in AA147-treated *ATF6*^*hom*^ retinal organoids compared to RP22-treated *ATF6*^*hom*^ organoids after 50 days of treatment. The violin plots show expression levels of 244 cone photoreceptor genes and 20 rod photoreceptor genes in AA147-treated *ATF6*^*hom*^ retinal organoids relative to RP22-treated *ATF6*^*hom*^ organoids. Thick dashed line marks the mean, and the thin horizontal lines delimit the upper and lower quartiles of gene expression. The complete gene sets are shown in Supplemental Table S6. ****p<0.0001, t=4.776. **(g)** The relative ratios of cone gene expression normalized to rod gene expression in *ATF6*^*hom*^ retinal organoids compared to *ATF6*^*het*^ organoids, and in AA147-treated compared to RP22-treated *ATF6*^*hom*^ retinal organoids are shown. ****p<0.0001.

Next, we examined the effects of AA147 in patient iPSC-derived retinal organoids expressing the Y567N ATF6 mutant. We modified our retinal organoid differentiation protocol to incorporate AA147 or the inactive analog RP22 to media retinal organoids from day 120 of differentiation, and then, further differentiated these organoids in the presence of compounds for another 50 days (**Fig. S4**). We then analyzed AA147- and RP22-treated organoids at day 170 (D170) of differentiation. We confirmed induction of ATF6 transcriptional targets in AA147-treated retinal organoids through increased levels of ATF6 transcriptional target genes as compared to RP22-treated organoids (**Fig. 4c, Supplemental Table 5**). When we microscopically examined the surface morphology of these retinal organoids at D170, we saw the emergence of ovoid protrusions on the surface of AA147-treated organoids, while RP22-treated organoids remained completely smooth (**Fig. 4d, 4e**). These protrusions on D170 AA147-treated *ATF6*^*hom*^ retinal organoids resembled smaller versions of the conical structures seen in the older D290 *ATF6*^*het*^ and *ATF6*^*+/+*^ retinal organoids (**Fig. 1, Fig. S2**), suggesting that AA147 restored cone photoreceptor development. To further define the impact of AA147 on mutant cones, we performed RNA-seq on pooled RNA isolated from the D170 organoids. Consistent with an AA147-induced rejuvenation of cone photoreceptors in *ATF6*^*hom*^ retinal organoids, we saw higher median expression of many cone transcripts, relative to rod transcripts, in *ATF6*^*hom*^ retinal organoids treated with AA147 as compared to RP22 treated *ATF6*^*hom*^ retinal organoids (**Fig. 4f, Supplemental Table 4, Supplemental Table 6**). Cone transcripts induced by AA147 included visual pigments and phototransduction proteins such as CNGB3 and OPN1S*W* (**Fig. S5**). When compared to *ATF6*^*het*^ retinal organoids, we found that *ATF6*^*hom*^ retinal organoids showed a ∼50% reduction in cone gene expression levels relative to rod genes, and AA147 treatment partially restored this deficit (**Fig. 4f, 4g**). These results demonstrate that our small molecule proteostasis strategy rescues the transcriptional activity of a human mutant ATF6 protein and stimulates growth of cone photoreceptors in patient retinal organoids carrying this Class 1 *ATF6* disease-causing variant.

## Discussion

ATF6 is a key transcriptional regulator of the UPR that ensures the ER organelle synthesizes high quality proteins and lipids in human cells throughout life. However, it is unknown why hypomorphic variants in *ATF6* cause vision loss diseases in people. Here, we combine human stem cell retinal organoid models, high resolution patient retinal imaging, and small molecule proteostasis modulators to identify cellular and molecular causes for vision loss arising from ATF6 inactivation. We identify absence of cone inner/outer segment structures in retinal organoids and patient retinas as an underlying cellular pathology for loss of vision in these patients. These cellular structures are essential for cones to detect photons and transduce electrical activity in photoreceptors in response to light. At the molecular level, loss of the entire cone phototransduction apparatus explains why patients carrying *ATF6* disease-causing variants have visual symptoms identical to those seen in patients carrying inactivating variants in cone phototransduction genes ^6^.

Our retinal organoid studies identify a novel molecular strategy to revive diseased cones in patients. In patients with variants that impede ATF6 exit from the ER, we show that we can overcome this defect by altering the equilibrium between ATF6 oligomers and monomers in the ER using a small molecule that positively targets the activation mechanism of ATF6. By favoring the monomeric, reduced ATF6 competent for downstream trafficking, we increase the amount of ATF6 protein that can escape from the ER. Using this strategy, we demonstrate that AA147 treatment restores ATF6 transcriptional activity of a Class 1 *ATF6* disease-causing variant and partially rejuvenates diseased cones in retinal organoids bearing this variant. We anticipate even greater cone rescue can be achieved through optimization of small molecule treatment regimens, and that this strategy will be effective for additional *ATF6* variants that cause vision loss.

**Figure S1.**
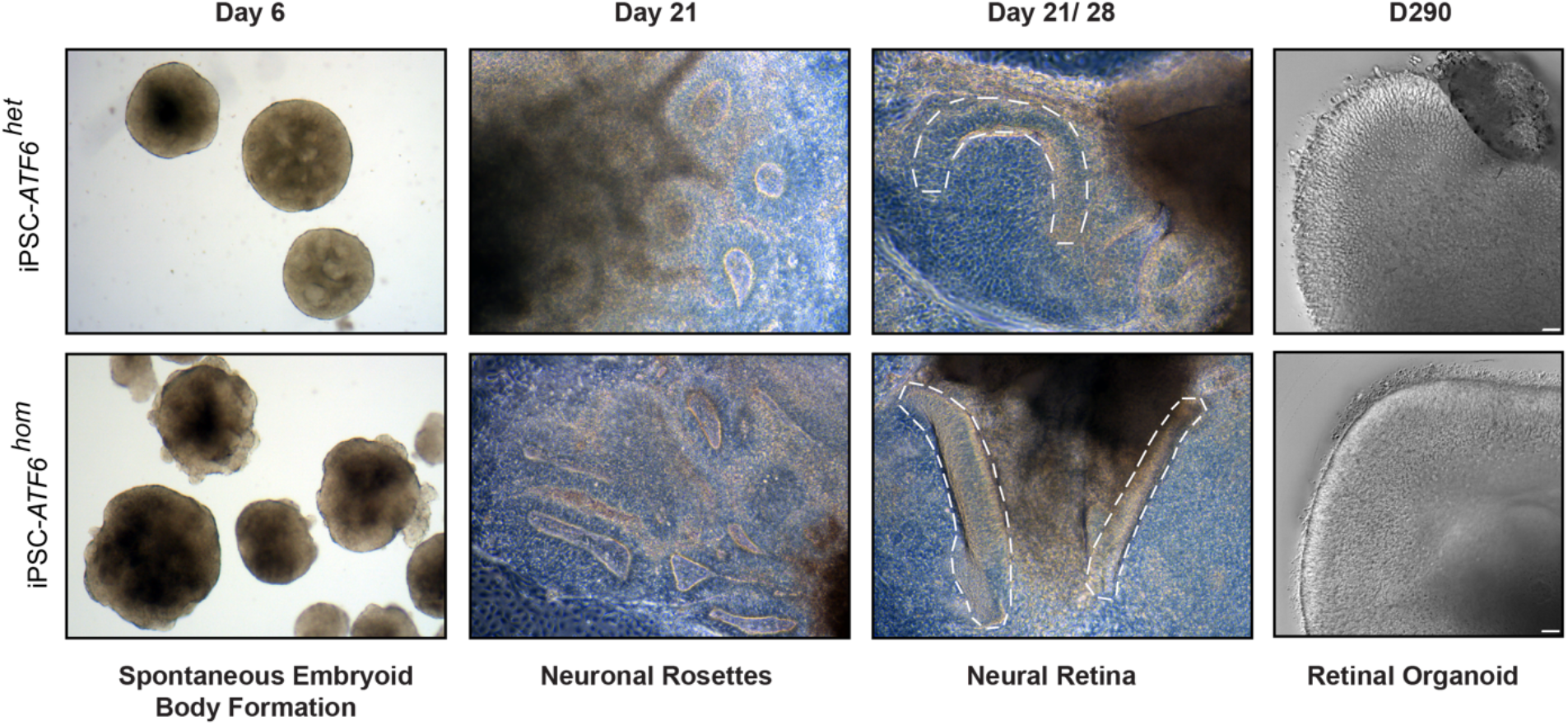
*ATF6* mutant iPSCs efficiently form embryoid bodies, neuronal rosettes, neural retina (highlighted in dashed circles), and retinal organoids. Representative photographs of brightfield images from D6-D28 and DIC image on D290 are shown of *ATF6*^*het*^ iPSCs (top row) or *ATF6*^*hom*^ iPSCs at the indicated timepoints of differentiation into retinal organoids.

**Figure. S2.**
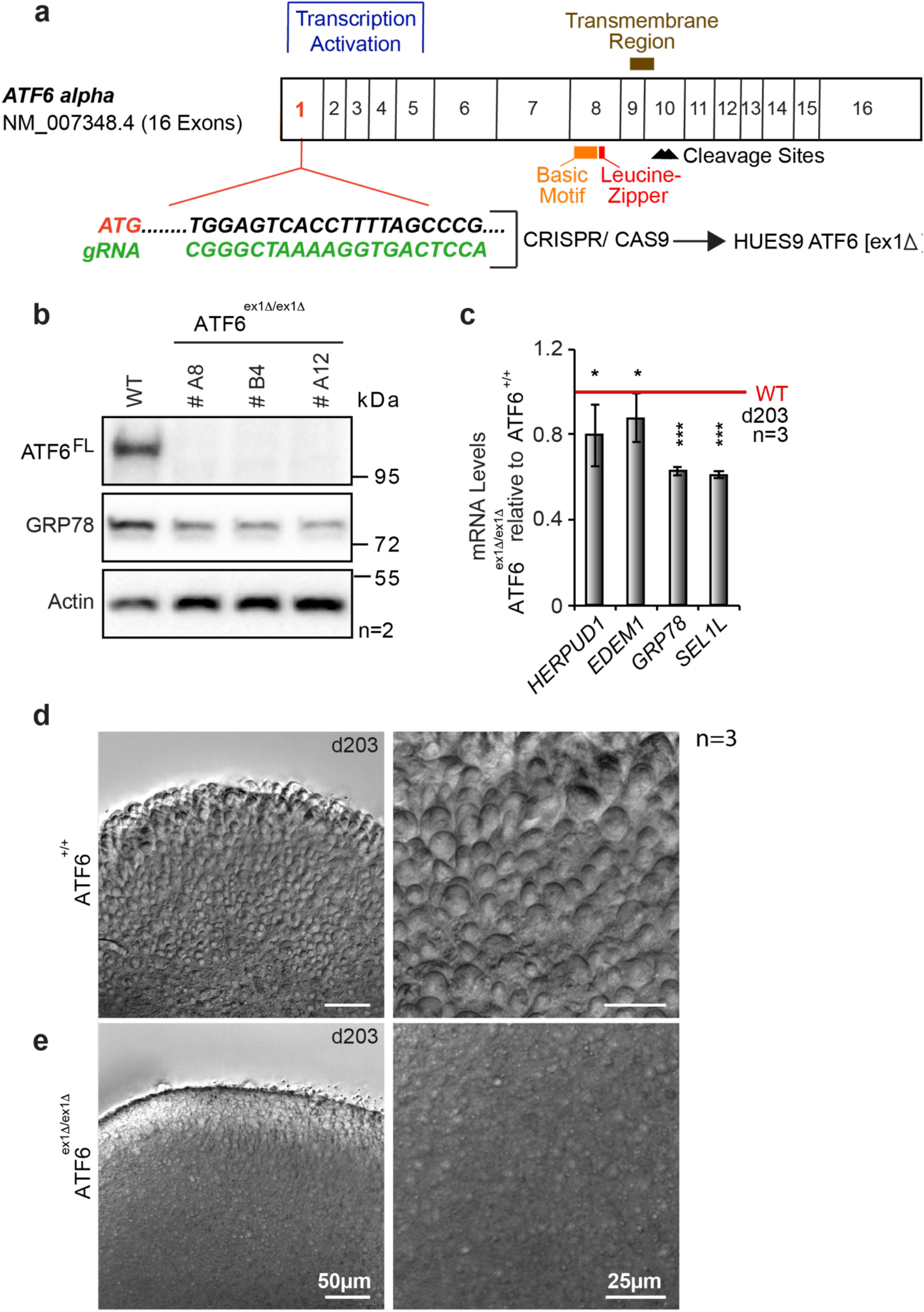
hESC *ATF6*^*ex1Δ/ex1Δ*^ retinal organoids do not form cone structures. **(A)** Schematic of CRISPR/Cas9 gene-editing strategy to generate *ATF6*^*ex1Δ/ex1Δ*^ exon 1 deletion of human ATF6 alpha hESCs. (**b)** Cell lysates from parental *ATF6*^*+/+*^ and several *ATF6*^*ex1Δ/ex1Δ*^ hESC clones were immunoblotted for ATF6, GRP78/BiP, and actin (loading control). (**c)** qPCR analysis shows reduced expression levels of ATF6 transcriptional targets, *HERPUD1, EDEM1, GRP78*, and *SEL1* in *ATF6*^*ex1Δ/ex1Δ*^ retinal organoids normalized to levels in parental *ATF6*^*+/+*^ retinal organoids at 203 days of differentiation (red line; mean +/- s.d. of n=3 independent retinal organoids; two-tail Student’s *t*-test; *<0.05, *** <0.005). (**d, e)** DIC microscopy of retinal organoids reveals abundant nascent cone inner/outer segment ovoid structures on the surfaces of hESC *ATF6*^*+/+*^ retinal organoids (**d**) and absence of cone structures in hESC *ATF6*^*ex1Δ/ex1Δ*^ retinal organoids at 203 days (**e**). Right panels show high power magnifications from left panel images. n=3.

**Fig. S3.**
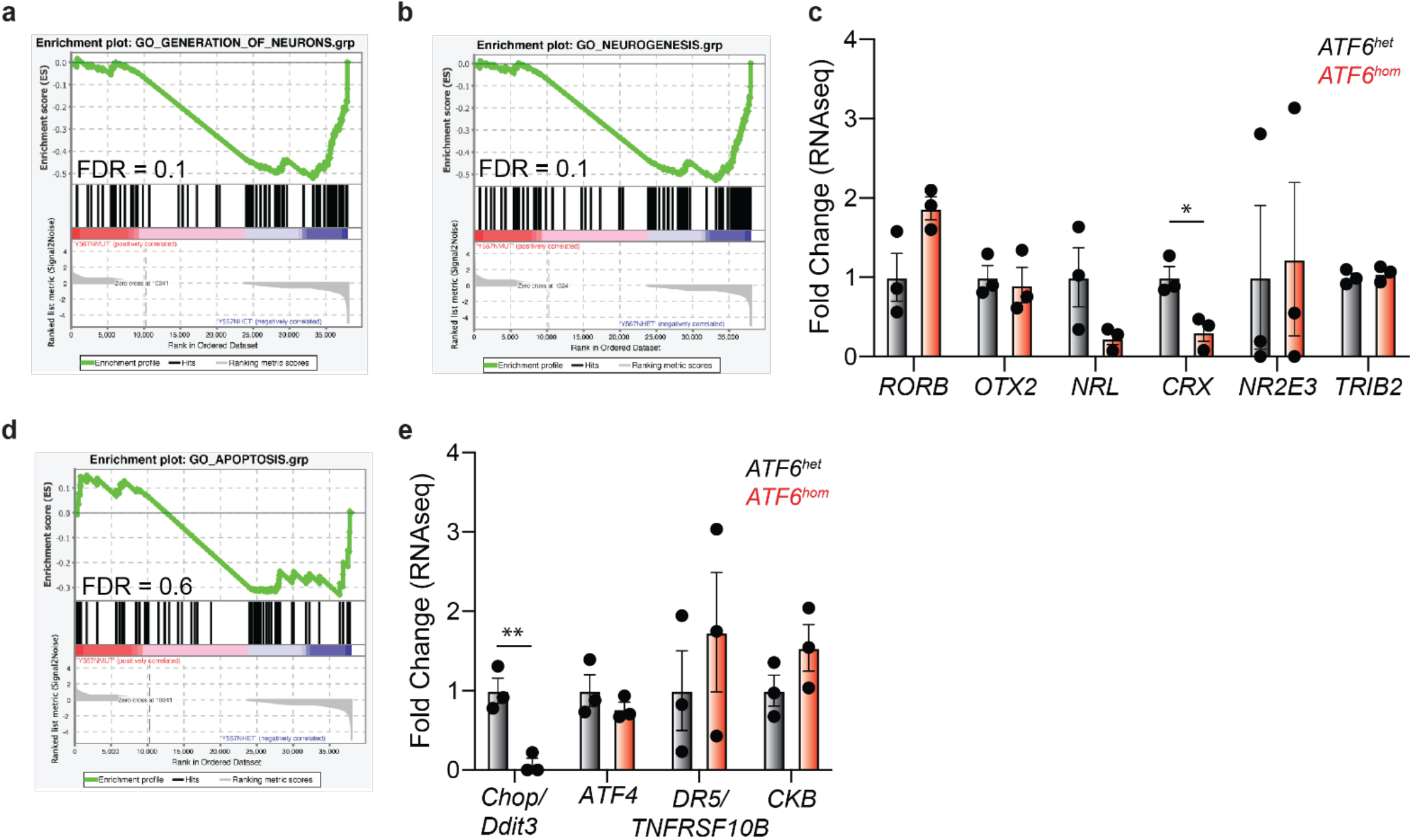
Retinal development and apoptosis pathways are not significantly altered in ATF6 mutant retinal organoids. (**a, b**) Gene set enrichment analysis (GSEA) plot of expression levels of neuronal generation and neurogenesis genes in *ATF6* mutant retinal organoids, relative to *ATF6*^*het*^ retinal organoids, with False Discovery Rate (FDR) as indicated. (**c**) Cone photoreceptor fate specification genes transcripts measured by RNA-seq from *ATF6*^*hom*^ retinal organoids are shown relative to transcript levels in *ATF6*^*het*^ retinal organoids. Error bars represent mean +/- standard deviation, and points represent individual retinal organoids. * p<0.05. (**d**) GSEA plot of expression levels of apoptosis genes in *ATF6* mutant retinal organoids, relative to *ATF6*^*het*^ retinal organoids. (**e**) ER stress-induced pro-apoptosis genes in *ATF6* mutant retinal organoids, relative to *ATF6*^*het*^ retinal organoids. Error bars represent mean +/- standard deviation, and points represent individual retinal organoids. ** p<0.01.

**Figure S4.**
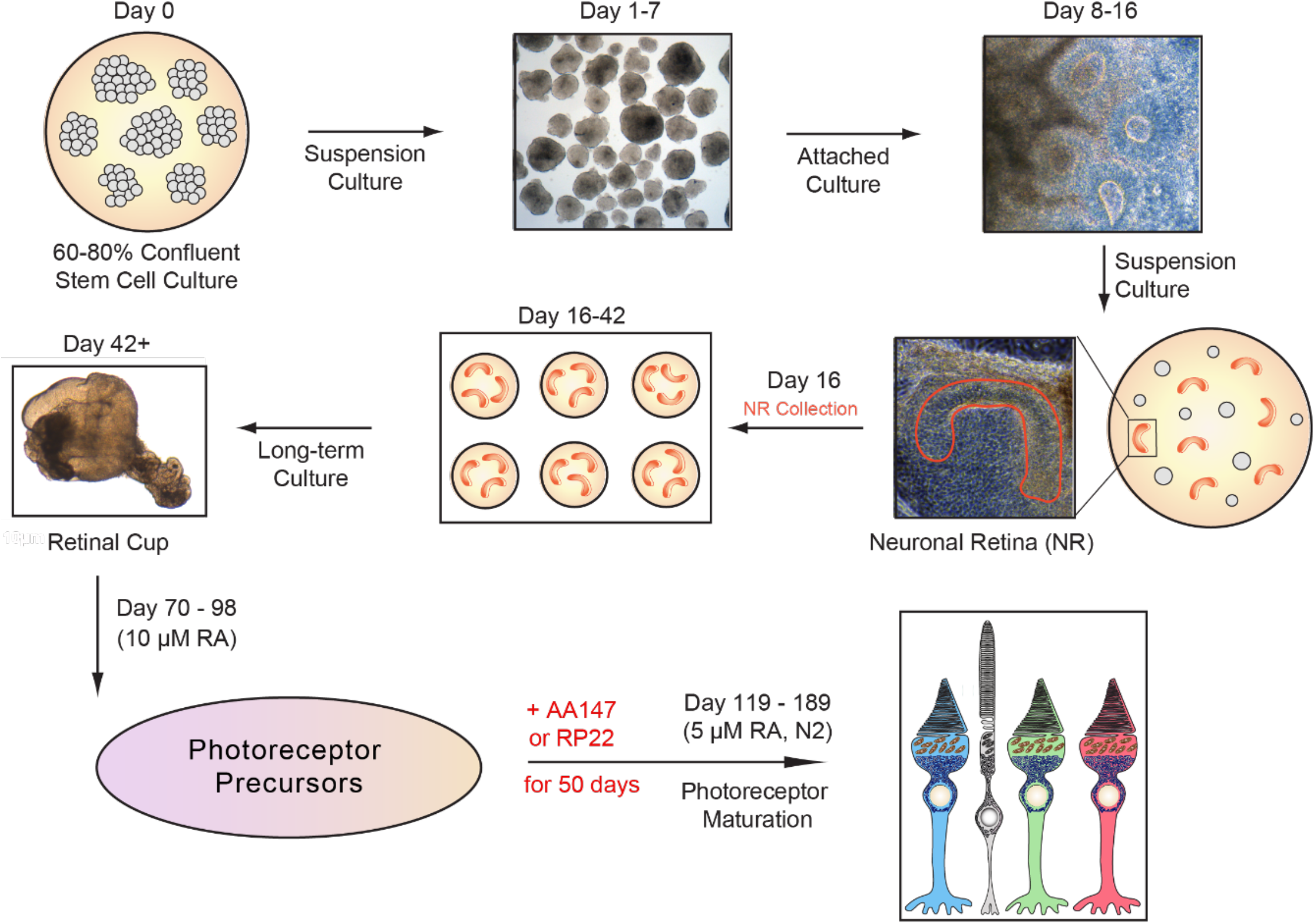
Schematic of protocol for small molecule proteostasis treatment of retinal organoids. Pluripotent iPSCs were differentiated into retinal organoids as illustrated and previously described.^9^ AA147 (10 μM) or RP22 (10 μM) was added to medium at day 120, fresh media with the small molecules was changed every 36h, and retinal organoids were cultured for 50 days during photoreceptor maturation. Retinal organoids were analyzed at day 170 of differentiation.

**Figure S5.**
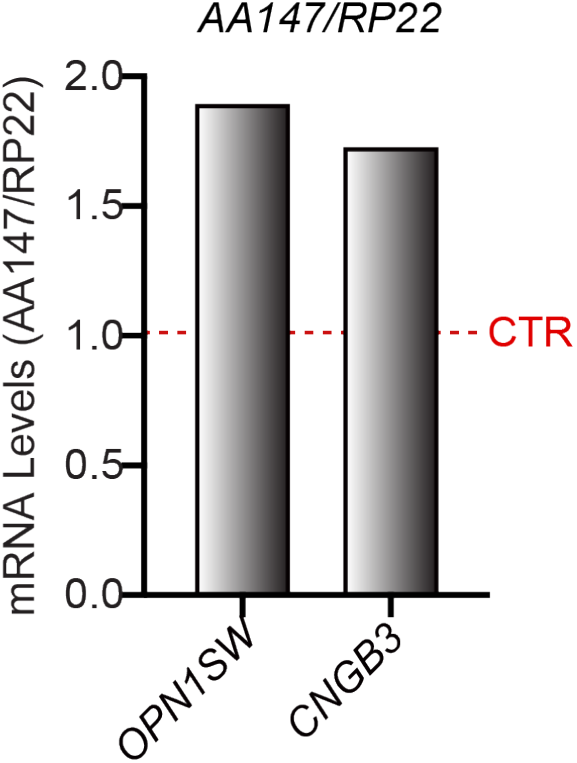
OPN1SW and CNGB3 transcripts detected by RNA-seq of AA147-treated *ATF6* mutant retinal organoids are shown relative to levels in RP22-treated *ATF6* mutant retinal organoids (CTR, dashed line). RNA was prepared from pooled replicates of two organoids per treatment condition from two independent experimental repeats (n=2); data were collected using three technical repeats (n=3).

## Materials and Methods

### Cell Culture

Human somatic cell lines (fibroblasts and HEK293 cells) were maintained at 37 °C, 5% CO2 in Dulbecco’s modified Eagle medium (Mediatech), supplemented with 10% FCS (Mediatech), and 1% penicillin/streptomycin (Invitrogen). Primary human fibroblast cells were established from skin biopsies of achromatopsia patients expressing two copies of *ATF6* mutant alleles (*ATF6*^*hom*^) or unaffected family members carrying a single *ATF6* mutant allele and a wild-type *ATF6* copy (*ATF6*^*he*t^), and these included Y567N, R324C, and D564G missense mutations as previously described ^6-8^. Fibroblasts were reprogrammed into iPSCs using the CytoTune-iPS 2.0 Sendai Reprogramming kit (Life Technologies), and 3 to 5 independent clones were created from each patient donor. All iPSC lines expressed pluripotency markers, had normal karyotypes, and were able to differentiate into all 3 germ layers using the embryoid body (EB) procedure. Early passage hESC H9 clones were obtained from the Human Embryonic Stem Cell Core Facility at the Sanford Consortium for Regenerative Medicine at the University of California, San Diego (UCSD). All iPSC and hESC lines were maintained on Corning Matrigel coated dishes (Corning Inc.) using mTESR1 medium (Stemcell Technologies) at 37°C and 5% CO2. Medium was changed daily, and pluripotent stem cells were passaged every 5 to 7 days in the ReLeSR medium (Stem Cell Technologies). Pluripotent iPSC and hESC were differentiated into retinal organoids as previously described (^9^ and **Fig S1, Fig S4**). The retinal organoid data presented in this study all derive from the passages 10-17 iPSCs carrying the Y567N ATF6 mutation.

CRISPR/Cas9 editing of the *ATF6* gene in hESCs was carried out by ThermoFisher Scientific Cell Model Services using the gRNA “cgggctaaaaggtgactcca” to introduce indels into exon 1 of the human *ATF6A* gene. Sanger and next-generation sequencing (NGS) of isolated and expanded hESC clones revealed homozygous knockout clones (*ATF6*^*exΔ1/exΔ1*^). NGS of the parental unedited hESCs (*ATF6*^*+/+*^) and three expanded knockout clones (*ATF6*^*exΔ1/exΔ1*^) further revealed no off-target cleavage in gene-edited clones compared to parental clone.

All stem cell studies followed ethical guidelines with ESCRO/IRB approval.

### Compounds

ATF6-activating compound AA147 and its analog RP22 were prepared in DMSO as stock solution at 10 mM, and working aliquots were prepared and stored at −20°C. The compound was used in cell culture medium at a final concentration of 10 μM. ER stress-inducing compound, DTT, (BioPioneer Inc.) was dissolved in water and added to the cell culture media at final concentration of 2 mM.

### Molecular Biology

Cells were lysed and total RNA collected using the RNeasy mini kit, according to manufacturer’s instructions (Qiagen). mRNA was used for RNA-seq analysis or prepared for qRT-PCR using the iScript cDNA Synthesis Kit (Bio-Rad). cDNA was used as template in SYBR green qPCR supermix (Bio-Rad). PCR Primers used include: human *RPL19*, 5′-ATGTATCACAGCCTGTACCTG-3′ and 5′-TTCTTGGTCTCTTCCTCCTTG-3′; human *BIP*/*GRP78*, 5′-GCCTGTATTTCTAGACCTGCC-3′ and 5′-TTCATCTTGCCAGCCAGTTG-3′; human *HERPUD1*, 5′-AACGGCATGTTTTGCATCTG-3′ and 5′-GGGGAAGAAAGGTTCCGAAG-3′; human *SEL1L*, 5′-ATCTCCAAAAGGCAGCAAGC-3′ and 5′-TGGGAGAGCCTTCCTCAGTC-3′; human *EDEM1*, 5′-TTCCCTCCTGGTGGAATTTG-3′ and 5′-AGGCCACTCTGCTTTCCAAC-3. *RPL19* mRNA levels served as internal normalization standards. qPCR condition was 95 °C for 5 min, 95 °C for 10 s, 60 °C for 10 s, 72 °C for 10 s, with 40 cycles of amplification.

### Immunoblotting Analysis

HEK293 cells expressing wild-type or mutant ATF6 were lysed in SDS lysis buffer (2% SDS in PBS containing protease and phosphatase inhibitors (Thermo Scientific). Protein concentrations of the total cell lysates were determined by BCA protein assay (Pierce). Equal amounts of protein were loaded onto 10% or 4–15% Mini-PROTEAN TGX precast gels (Bio-Rad) and analyzed by Western blot. The following antibodies and dilutions were used: anti-FLAG at 1:5,000 (Sigma-Aldrich); anti-BiP/GRP78 (GeneTex) at 1:1,000, and anti-GAPDH (Santa Cruz Biotech) at 1:5,000. After overnight incubation with primary antibody, membranes were washed in TBS with 0.1% Tween-20, followed by incubation of a horseradish peroxidase-coupled secondary antibody (Cell Signaling). Immunoreactivity was detected using the SuperSignal West chemiluminescent substrate (Pierce). GAPDH levels were assessed as a loading control as indicated.

### RNA-seq Analysis

RNA-seq was performed as previously described ^30^. Briefly, RNA was isolated from individual D290 retinal organoids or pooled D170 retinal organoids using the Qiagen RNAeasy Mini Kit according to the manufacturer’s instructions. RNA sequencing was performed by BGI Americas on the BGI proprietary platform (DNBseq), providing single-end 50 bp reads at 20 million reads per sample. Alignment of the sequencing data was performed using DNAstar Lasergene SeqManPro to the GRCh37.13 human genome reference assembly. Assembly data were then imported into ArrayStar 12.2 with QSeq (DNAStar Inc) to quantify the gene expression and normalized reads per kilobase million (RPKM). Differential expression analysis and statistical significance calculations between conditions was assessed using DESeq in R using a standard negative binomial fit for the aligned counts data and are described relative to the indicated control (Supplementary Tables S1, S4). Gene Ontology (GO) analysis was performed using Panther (geneontology.org; Supplementary Table S3). Geneset enrichment analysis (GSEA) was performed using denoted genesets from GO on the GSEA platform (gsea-msigdb.org) ^31,32^. Violin plots comparing mutant and control retinal organoids were generated using the fold change data from the differential expression analysis for human rod and cone genes ^19^ (Supplementary Table S2). Violin plots demonstrating changes in UPR-associated gene expression were generated using the fold change data from differential expression analysis for described genesets of UPR transcriptional targets ^30^(Supplementary Table S5). Violin plots comparing mutant retinal organoids treated with AA147 (10 µM) or the inactive analog, RP22 (10 µM) were generated using the normalized fold change data from the differential expression analysis for rod and cone genes, where fold change values of all genes of interest were normalized to the mean fold change of rod genes (Supplementary Table S6). Data used for generating violin plots were subject to ROUT outlier testing in GraphPad Prism.

### Immunofluorescence and Confocal Microscopy

Retinal organoids were washed three times in 1x PBS (w/o Mg^2+/^Ca^2+^) for 5 min, prior to fixing in 4% PFA for 45 min and gentle agitation at room temperature. After fixing, organoids were washed three times in 1x PBS (w/o Mg^2+^/Ca^2+^) for 5 min, followed by 30% (w/v) sucrose treatment for 48 hrs to allow organoids to settle down. Organoids were embedded in OCT and cryosectioned in 10 µm sections (Leica CM 1950), slides were stored at −20C. Prepared slides were adjusted to room temperature, washed three times in 1x PBS (w/o Mg^2+^/Ca^2+^) for 5 min. Samples were blocked and permeabilized for at least one hour in blocking buffer containing 1% (w/v) BSA/ 1x PBS (w/o Mg^2+^/Ca^2+^), 0.1% Triton X-100, and 5% goat serum. Primary antibody was prepared in blocking solution and incubated overnight at 4C with gentle agitation. Primary antibodies used; rabbit anti-GNAT1 (Sigma), mouse anti-rhodopsin (Santa Cruz Biotechnologies) and rabbit anti-cone opsin (OPN1MW/ LW; EMD Millipore). Slides were the washed six times in 1x PBS (w/o Mg^2+^/Ca^2+^) with 0.1% Triton X-100 for 5 min at room temperature with gentle agitation. Secondary antibody was prepared in blocking buffer and incubated for an hour at room temperature. Secondary antibodies included Alexa546 goat anti-mouse (red) antibody (Molecular Probes) and Alexa488 goat anti-rabbit (green) antibody (Molecular Probes) used at 1:250 dilution. Slides were then washed six times in 1x PBS (w/o Mg^2+^/Ca^2+^) for 5 min at room temperature with gentle agitation followed by mounting samples with ProLong Gold antifade reagent with DAPI (Invitrogen). Images were collected with an Olympus FluoView-1000 confocal microscope and processed using Olympus FluoView Ver.2.0a Viewer software at University of California, San Diego, School of Medicine Microscopy Core.

### Live Organoid Imaging and Video Preparation

Organoids were placed in a chamber slide (Ibidi) without a lid and mounted in a stage top incubator (H301-K-FRAME with Koehler Lid, Okolab). The temperature (37 °C), CO2 (5 %) and humidity (90 %) was controlled by a H301-T-UNIT-BL-PLUS stage incubation system (Okolab), CO2-UNIT-BL gas controller (Okolab) and HM-ACTIVE Humidity Controller (Okolab). All images were collected on an Eclipse Ti2 inverted microscope (Nikon Instruments) equipped with a Plan Apo λ 20x NA 0.75 (Nikon Instruments). The DIC mode used two polarizers, two Nomarski prims (N1 and 20x, Nikon Instruments) and diascopic LED illumination. Images were acquired with a monochrome DS-Qi2 sCMOS camera (Nikon Instruments) controlled by NIS-elements software (Nikon Instruments). Z-series optical sections were collected with a step-size of 0.9 µm, using the Ti2 ZDrive. The Z-series were processed using local intensity algorithm (25 °, radius = 5.46 µm), the focused image (balanced Z-map method) of the Z-series is displayed using NIS-Elements software (Nikon Instruments). To generate videos, a montage of the processed Z-series with 10 copies of the focused image is played back at 20 frames per second.

### Fixed Retinal Organoid Imaging

Whole organoids were stained using the same protocol as described for OCT embedded sections. After staining, the entire organoid was placed in a 35 mm round tissue culture dish without lid (FD35-100, World Precision Instruments). To stabilize whole organoids during images, organoids were maintained NIM III media supplemented with 50% glycerol. All images were collected with an X-Light V2 LOV (Crestoptics, S.p.A.) with 50 µm pinholes spinning on an Eclipse Ti2 inverted microscope (Nikon Instruments) equipped with a Plan Apo VC 60xA WI NA 1.2 objective lens. The DIC mode used two polarizers, two Nomarski prims (N2 and 60xII, Nikon Instruments) and diascopic LED illumination. Cone opsin (OPN1MW/LW, EMD Millipore) was detected using Alexa Fluor 488 and was excited with the 473 nm line from a Celesta light engine (Lumencor), Rod opsin (Santa Cruz Biotechnologies) was detected using Alexa Fluor 546 and was excited with the 545 nm line from a Celesta light engine (lumencor). The emission was collected using a penta dichroic mirror (FF421/491/567/659/776-Di01, Semrock) and a penta emission filter (FF01-441/511/593/684/817, Semrock). Images were acquired with an Iris 15 CMOS camera (Photometrics) controlled by NIS-Elements software (Nikon Instruments). Z-series optical sections were collected with a step-size of 0.3 µm, using the Ti2 Z-Drive. The Z series is processed with Fiji ^33^. For the DIC Z-series, an FFT bandpass filter (filtering large structures down to 40 pixels, small structures up to 2 pixels, and tolerance of direction 5%) was applied followed by an unsharp mask (with radius of 5 pixels and mask weight 0.60). For the green and red fluorescence Z-series, an FFT bandpass filter (filtering large structures down to 400 pixels, small structures up to 4 pixels, and tolerance of direction 5%) was applied followed by a background subtraction using a rolling ball (radius of 50 pixels). A 3D image and movie were created using the Volume View and Movie Maker of the NIS-Elements Analysis software (Nikon Instruments).

### AOSLO Retinal Imaging

Informed consent was obtained from a patient with confirmed *ATF6* disease-causing variants and a normal control. Prior to imaging, the combination of tropicamide (1%) and phenylephrine hydrochloride (2.5%) was used for cycloplegia and pupillary dilation. AOLSO videos were acquired and processed as previously described ^18,34^. This study followed the tenets of the Declaration of Helsinki and was approved by the institutional review board at the Medical College of Wisconsin.

### Statistical analysis

Student two-tailed t tests (for paired samples) were performed to determine P values. A value of P < 0.05 was considered significant. **P < 0.05 and ***P < 0.005. For RNA-Seq, statistical significance of differences in fold change expression for rod versus cone genes was calculated using two-tailed Student’ t-test. ****P<0.0001. Genesets comprising different stress-responsive signaling pathways were selected as previously described ^30^. Statistical significance of differences in fold change expression values for stress-responsive genesets were calculated using one-way ANOVA, and the reported adjusted P-value comparing the ATF6 geneset to the PERK/ATF4 geneset ^30^.

## Supporting information

SI Guide

## Acknowledgments

We thank Eric Griffis of the Nikon Imaging Center at UC San Diego and Elise Heon, Yang Hu, Ajoy Vincent, Doug Vollrath, and Sui Wang for helpful feedback and sharing materials.

## Funding

This work was supported by NIH grants AG046495, AG063489, EY027335, NS088485, P30NS047101; VA Merit awards I01BX002284, I01RX002340; California Institute for Regenerative Medicine DISC2-10973; and by grants from the National Institute for Health Research Biomedical Research Centre at Moorfields Eye Hospital NHS Foundation Trust and UCL Institute of Ophthalmology.

## Author contributions

H.K. and J.H.L. designed the project. H.K., D.B., W.-C.C., J.O., and N.J.G. performed the experiments. J.M.D.G., E.P., J.W.K., and R.L.W. analyzed the RNA-seq data. R.M., J.C. and M.M., analyzed the patient retinal imaging. H.K. and J.H.L. wrote the manuscript.

## Competing interests

J.K. declares that he is a board member and shareholder of Proteostasis Therapeutics Inc., Protego BioPharma, and Yumanity, which may develop ATF6 activators to treat degenerative diseases, although not for stem cell–associated purposes at this time. J.W.K. and R.L.W. are inventors on a patent describing the ATF6 activating compound used in this study that has been licensed to Protego BioPharma. All other authors declare they have no competing interests.

## Data and materials availability

The RNA-seq data have been deposited to the public National Center for Biotechnology Information GEO repository under the data identifier GSE106847.

## Notes

### Competing Interest Statement

J.W.K. declares that he is a board member and shareholder of Proteostasis Therapeutics Inc., Protego BioPharma, and Yumanity, which may develop ATF6 activators to treat degenerative diseases, although not for stem cell associated purposes at this time. J.W.K. and R.L.W. are inventors on a patent describing the ATF6 activating compound used in this study that has been licensed to Protego BioPharma. All other authors declare they have no competing interests.

