## Supplementary material for "The Endoplasmic Reticulum Proteostasis Regulator ATF6 is Essential for Human Cone Photoreceptor Development": SI Guide

| Parent Figure or Table | Filename<br><br>This should be the name the file is saved as when it is uploaded to our system, and should include the file extension. i.e.: *Smith_SourceData_Fig1.xls,* or *Smith_ Unmodified_Gels_Fig1.pdf* | Data description<br><br>i.e.: Unprocessed Western Blots and/or gels, Statistical Source Data, etc. |
|---|---|---|
| Fig. 3a | Table S1.xlsx | Normalized aligned counts and differential expression (DESeq) analysis of *ATF6*$^{hom}$ versus *ATF6*$^{het}$ retinal organoids |
| Fig. 3a, b, c | Table S2.xlsx | Fold change expression values for cone and rod genes of interest in ATF6[Y567N]$^{hom}$ retinal organoids |
| Fig. 3e, S3d | Table S3.xlsx | Gene ontology analysis of significantly upregulated and downregulated genes in ATF6[Y567N]$^{hom}$ retinal organoids |
| Fig. 4f | Table S4.xlsx | Differential expression (DESeq) analysis of *ATF6[Y567N]*$^{hom}$ organoids treated with AA147 versus RP22 |
| Fig. 4c | Table S5.xlsx | Fold change expression values for UPR target genes of interest in *ATF6[Y567N]*$^{hom}$ retinal organoids treated with AA147 |
| Fig. 4f | Table S6.xlsx | Fold change expression values for cone and rod genes, normalized to mean rod gene fold change, in *ATF6[Y567N]*$^{hom}$ retinal organoids and *ATF6[Y567N]*$^{hom}$ retinal organoids treated with AA147. |
| Fig. 1a, b | Supp Data Video 1 | Fixed Retinal Organoid Surface Analysis: Immunolabeling of fixed retinal organoid derived from *ATF6*$^{het}$iPSCs (green staining presents red/ green opsin; red staining presents rhodopsin). Prolonged *in vitro* culturing results in round ovoid protrusions emerged from the surfaces of *ATF6*$^{het}$ retinal organoids that were morphologically consistent |

| | | |
|---|---|---|
| | | with nascent cone inner/outer segments and contained cone opsin proteins. |
| Fig. 1c | Supp Data Video 2 | Live Retinal Organoid Surface Analysis: Cone structures on retinal organoid surfaces of *ATF6*$^{het}$ iPSCs generated from family members with normal vision |
| Fig. 1d | Supp Data Video 3 | Live Retinal Organoid Surface Analysis: Microscopic examination of the surfaces of *ATF6*$^{hom}$ patient retinal organoids showed no ovoid cone structures and instead revealed smoother contours consistent with fine packing of slender rods |
| Fig. S2d | Supp Data Video 4 | Live Retinal Organoid Surface Analysis of isogenic *ATF6*$^{+/+}$ retinal organoids: abundant ovoid cone structures on organoid surface are visible. but no ovoid protrusions on retinal organoids lacking ATF6, similar to what we observed in patient iPSC-derived retinal organoids |
| Fig. S2e | Supp Data Video 5 | Live Retinal Organoid Surface Analysis of isogenic *ATF6*$^{ex1\Delta/ex1\Delta}$ retinal organoids: no ovoid protrusions on surface of retinal organoids lacking ATF6. |
